# Nanobead-based single-molecule pulldown for single cells

**DOI:** 10.1101/2023.07.03.546659

**Authors:** Qirui Zhao, Yusheng Shen, Xiaofen Li, Yulin Li, Fang Tian, Xiaojie Yu, Zhengzhao Liu, Rongbiao Tong, Hyokeun Park, Levent Yobas, Pingbo Huang

## Abstract

Investigation of cell-to-cell variability holds critical physiological and clinical implications. Thus, numerous new techniques have been developed for studying cell-to-cell variability, and these single-cell techniques can also be used to investigate rare cells. Moreover, for studying protein-protein interactions (PPIs) in single cells, several techniques have been developed based on the principle of the single-molecule pulldown (SiMPull) assay. However, the applicability of these single-cell SiMPull (sc-SiMPull) techniques is limited because of their high technical barrier and special requirements for target cells and molecules. Here, we report a highly innovative nanobead-based approach for sc-SiMPull that is based on our recently developed microbead-based, improved version of SiMPull for cell populations. In our sc-SiMPull method, single cells are captured in microwells and lysed in situ, after which commercially available, pre-surface-functionalized magnetic nanobeads are placed in the microwells to specifically capture proteins of interest together with their binding partners from cell extracts; subsequently, the PPIs are examined under a microscope at the single-molecule level. Relative to previously published methods, nanobead-based sc-SiMPull is considerably faster, easier to use, more reproducible, and more versatile for distinct cell types and protein molecules, and yet provides similar sensitivity and signal-to-background ratio. These crucial features should enable universal application of our method to the study of PPIs in single cells.

**Statement of Significance:** Heterogeneity between single cells holds critical physiological and clinical implications. Characterization of protein-protein interactions (PPIs) and identification of the interacting partners of a specific protein are critical for elucidating the function and regulation of the protein. However, the applicability of the currently available techniques for studying PPIs in single cells is limited by their high technical barrier and special requirements for cell/proteins types. Our single-cell single-molecule pulldown (sc-SiMPull) assay in this study is not only substantially simpler and faster than existing sc-SiMPull methods, but also considerably more widely applicable—to all cell types and to both soluble and transmembrane proteins. These two crucial features should enable universal application of our method to the study of PPIs in single cells.

## Introduction

Heterogeneity between single cells is observed in not only tissues and organisms, but also populations of genetically identical monoclonal cells, and the study of cell-to-cell variability has attracted considerable attention recently; this is because in addition to revealing previously unknown regulatory mechanisms, the investigation can identify potential barriers for effective therapeutic intervention ^1–3^. For example, cell-to-cell variation among cancer cells leads to the expression of distinct surface receptors and could thereby cause the failure of targeted therapies that rely on the surface receptors as biomarkers, and cancer heterogeneity also influences dug resistance, which is the most challenging hurdle in oncology ^4^. Because conventional approaches used for studying cell populations obscure cell-to-cell variability, numerous new techniques have been developed—owing to notable advances in analytical methods and microfluidic tools—for studying single cells and cellular heterogeneity. Besides being used to study cell-to-cell variability, these single-cell techniques can be employed to investigate rare cells such as auditory hair cells, circulating tumor cells, stem cells, and a subset of immune cells ^5, 6^. Mature techniques have now been developed for genomic and transcriptional analyses in single cells, and techniques for protein analysis in single cells are emerging as well, including single-cell western blotting ^7^, single-cell secretion assay ^3^, tunable single-cell extraction ^8^, and single-cell mass spectrometric analysis ^2^.

A protein never works solo and invariably functions together with its regulatory proteins, auxiliary subunits, or effector proteins, and this typically requires physical interaction between the proteins. Therefore, these protein-protein interactions (PPIs) are essential in nearly all aspects of diverse cellular processes, and identification and validation of the interacting partners of a specific protein are critical for elucidating the function and regulation of the protein. For studying PPIs in single cells, several techniques have been reported ^9–11^, and because these methods are based on the principle of the single-molecule pulldown (SiMPull) developed by Jain et al. ^12^, the methods are referred to as single-cell SiMPull (sc-SiMPull) techniques. However, the applicability of these sc-SiMPull techniques remains limited because (1) their technical barrier is considerably high and thus the methods cannot be used in common biological laboratories ^9–11;^ (2) the techniques can be used for analyzing slowly diffusing and soluble molecules only in bacterial cultures ^10^ or adherent-cell cultures ^9^ but not in primary- or suspension-culture cells (such as blood cells and circulating tumor cells); or (3) the throughput of the technique is extremely low (designed for single zygotes) ^11^.

Here, we report a highly innovative nanobead-based approach for sc-SiMPull that is based on our recently developed microbead-based, improved version of SiMPull for cell populations ^6^. We believe that our nanobead-based method is not only substantially simpler and faster than existing sc-SiMPull methods, but also considerably more widely applicable—to all cell types and to both soluble and transmembrane proteins. These two crucial features should enable universal application of our method to the study of PPIs in single cells.

## Methods and Materials

### Materials

The following commercially available reagents were used: methanol (BDH1135), acetone (BDH1101), and 2-propanol (BDH1133), BDH Chemicals; Tris-HCl (Cat.# BP153-1), Fisher Scientific; sodium deoxycholate (Cat.# D6750), biotin-Alexa 488 (Cat.# 30574), and bovine serum albumin (BSA; Cat.# A7030), Sigma-Aldrich; NeutrAvidin (Cat.# 31000), Pierce; NP-40 (Cat.# N3500), United States Biological; PEI 25000 (Cat.# 23966-1), Polysciences; and PBS (Cat.# 10010-023), Gibco.

The other materials used were Quartz slides (Cat.# 7101, Sail Brand), coverslips (24 × 24 mm; Cat.# 48393230, VWR International), magnetic beads (diameter 70–130 nm; SV0100, Ocean Nanotech), and a permanent magnet (Cat.# N35, Hongshi Inc., China).

### Antibodies

These antibodies against proteins and affinity tags were from commercial sources: mouse monoclonal anti-HA (MMS-101p, Covance), mouse anti-FLAG (clone M2, Sigma-Aldrich), biotinylated goat anti-rabbit IgG (Cat.# 65-6140, Thermo Fisher), Alexa Fluor 488-conjugated goat anti-rabbit IgG (ab181448, Abcam), Alexa Fluor 647-conjugated goat anti-rabbit IgG (ab150079, Abcam), Alexa Fluor 647-conjugated goat anti-mouse IgG (ab150115, Abcam), and Alexa Fluor 568-conjugated goat anti-mouse IgG (ab175473, Abcam).

Rabbit anti-green fluorescent protein (GFP) serum was homemade by immunizing rabbits housed in the Animal Care Facility at the Hong Kong University of Science and Technology.

### Solutions

T50-BSA buffer contained 50 mM NaCl, 0.1 mg/mL BSA, 10 mM Tris-HCl, pH 8.0; T50-Tween 20 buffer contained 50 mM NaCl, 0.1% Tween 20, 10 mM Tris-HCl, pH 8.0; both solutions can be stored at 4°C for up to 1 month. The lysis buffer contained 150 mM NaCl, 1 mM EDTA, 1% v/v NP-40, 10 mM Tris, protease inhibitors (cOmplete Mini, Roche), pH 7.5 adjusted with HCl; we recommended preparing fresh lysis buffer for each use.

### Fabrication of microwell arrays for cell-trapping

Cell-trapping microwell arrays were fabricated by generating microwell-patterned PDMS membranes on a glass coverslip. Briefly, a silicon mold featuring micropillar patterns was first fabricated using the Bosch deep-reactive ion etching process. Pillar diameter (15–50 μm) and interspacing (4 × pillar diameter, edge to edge) were controlled using standard photolithographic techniques, with the etching depth of the pillars being controlled at 75 μm. Next, the fabricated silicon mold was diced into chips sized 1.0 × 1.0 cm_2_ and silanized using dimethyldichlorosilane (Sigma-Aldrich) vapor in a vacuum chamber overnight, after which the mold was sequentially washed with acetone and deionized water and dried using N^2^ before use. Subsequently, 5 μL of pre-degassed PDMS pre-polymer (10:1, weight ratio of base to curing agent) was gently pipetted onto the silanized silicon mold and a piece of glass coverslip was brought in close contact with the chip to allow uniform spreading of the pre-polymer across the entire chip. The coverslip was pretreated with oxygen plasma and washed with acetone, 2-propanol, and deionized water in a tabletop ultrasonic cleaner (Bransonic), and then a 100 g weight was placed over the coverslip to ensure that the microfabricated pillars completely penetrated the pre-polymer layer and touched the coverslip. The entire assembly was placed in a 60°C oven for 3 h, and then after removing the weight, the coverslip together with the microwell-patterned PDMS membrane was slowly peeled off from the mold, which formed the cell-trapping device.

### Cell culture and transfection

Cells were cultured and transfected as previously described ^13^. HEK293T cells (RRID: CVCL_1926) were obtained from ATCC; the cells were assumed to be authenticated by ATCC and were not further authenticated in this study. The cell line routinely tested negative for mycoplasma contamination and was maintained in Dulbecco’s modified Eagle medium (DMEM) supplemented with 10% fetal bovine serum and 100 U/mL penicillin/streptomycin (Life Technologies) in an atmosphere of 95% air-5% CO_2_ at 37°C. Cells were transfected using PEI (3 μL/μg plasmid).

### Preparation of primary antibody-coated nanobeads

All procedures were performed at room temperature. First, 10 μL of streptavidin-coated magnetic nanobeads were mixed with 100 μL of 10 nM biotinylated secondary antibody in PBS in an Eppendorf tube, and after incubation for 10 min, the beads were pulled down using a permanent magnet, washed once with PBS, and incubated for 20 min with a primary antibody against the bait protein in PBS-BSA buffer (PBS containing 10 mg/mL BSA). The nanobeads were washed again with PBS and then stored in 100 μL of PBS until use. We recommended using the beads within 2 h at room temperature.

### Protein pulldown from cell populations by using magnetic nanobeads

Approximately 10^6^ cells were lysed using 100 μL of lysis buffer in an Eppendorf tube and the lysates were then mixed with 30 μL of primary antibody-coated nanobeads. After incubation for 30 min, the nanobeads were washed with 200 μL of wash buffer thrice, resuspended in a final volume of 5 μL of PBS, and transferred onto a glass slide and covered with a coverslip (18 × 18 mm). The coverslip was precleaned by sonicating in acetone, isopropanol, and water (5 min each). The 5 μL of nanobead-containing PBS was effectively contained between the coverslip and slide due to the capillary effect (also see Fig. 1).

**Figure 1.**
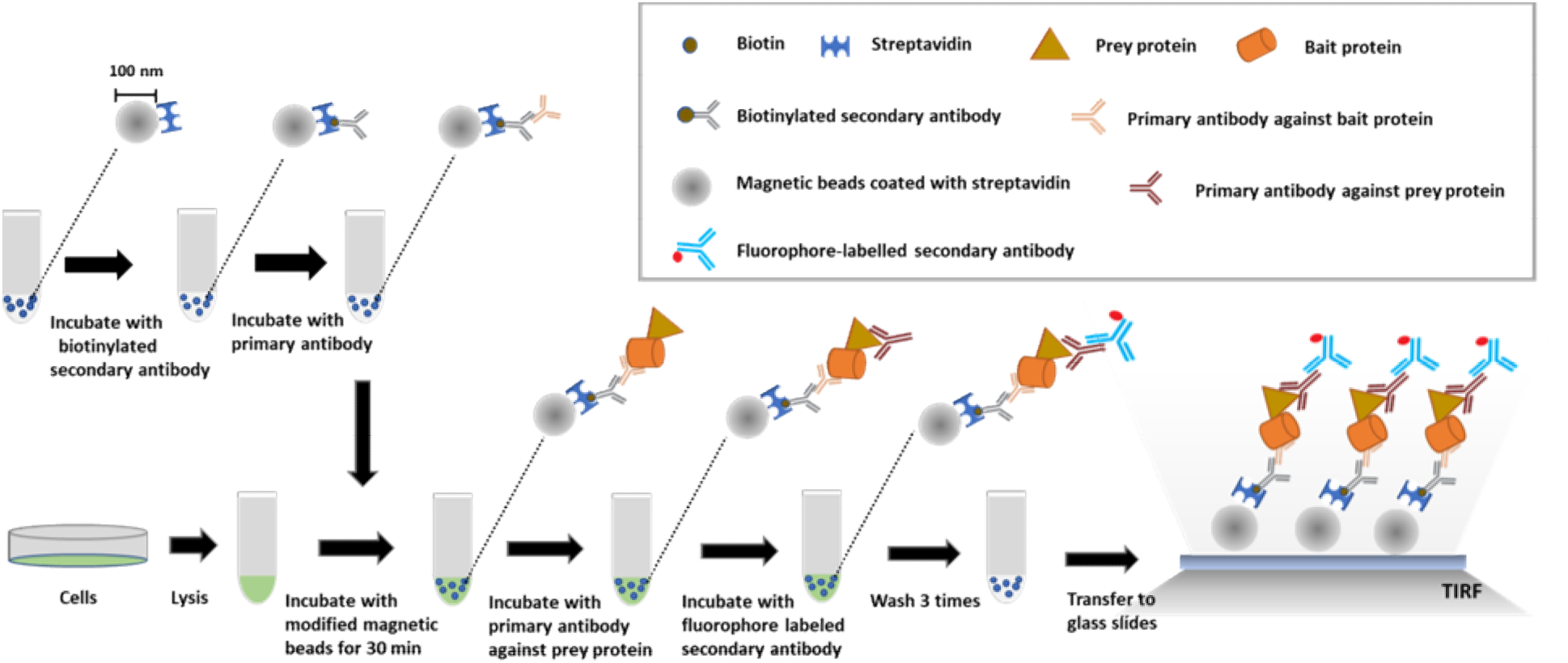
Schematic of nanobead-based SiMPull for cell populations. For capturing the target (bait) protein, streptavidin-coated magnetic nanobeads are modified by immobilizing a biotinylated 2^nd^ antibody and a bait-specific 1^st^ antibody on the nanobead surface. Cells are lysed and the magnetic nanobeads are added to the lysates and incubated for 30 min in an Eppendorf tube. After capture by the magnetic nanobeads, the prey protein is detected using a specific 1^st^ antibody and a fluorescently labeled 2^nd^ antibody. The nanobeads are washed thrice, transferred to a glass slide, covered with a coverslip, and imaged using a TIRF or confocal microscope. Some of the washing steps are omitted in the schematic for the sake of simplicity (see additional details in “Methods and Materials”), and between solution changes, the nanobeads are pulled down and held by applying a magnetic force. If the bait protein or the prey protein is fluorescently labeled, the protein can be visualized directly without immunostaining.

In the case of fluorescently tagged proteins, the proteins captured on the nanobeads could be imaged immediately under a total internal reflection fluorescence (TIRF) microscope or a confocal microscope. However, when the prey protein was not fluorescently tagged, 100 μL of 10 nM primary antibody against the prey protein in T50-BSA buffer was added into the Eppendorf tubes containing the samples, and after incubation for 20 min, the beads were washed with 200 μL of wash buffer thrice and incubated for 15 min with 50 μL of 20 nM fluorophore-labeled 2^nd^ antibody in T50-BSA buffer; lastly, after washing thrice more with the wash buffer to remove unbound antibodies, the nanobeads were resuspended in a final volume of 5 μL of PBS and imaged under a microscope.

### Cell-trapping for studying single cells

To the surface of the aforementioned microwell array chips, 300 μL of suspended cells in PBS (∼ 10^6^ cells/mL) was applied and the cells were allowed to settle by gravity for 5–10 min (depending on the cell size), a process that could be monitored under a microscope. After single-cell occupancy reached 30% of total microwells, the cell suspension was removed from the chip surface by pipetting and the chip was washed thrice with PBS to obtain a clean background.

### Protein pulldown by magnetic nanobeads in microwells

To the surface of chips harboring trapped cells, 100 μL of antibody-coated magnetic nanobeads in PBS was evenly applied and the chip was placed on a permanent magnet (∼800 Gs on surface) for 3 min to draw ∼80% of the nanobeads down to the chip surface and the bottom of the wells (Supplementary Fig. 7). After removing excess PBS from the chip surface by using a piece of Kimtech paper, 0.5 mL of lysis buffer was added gently to the chip surface from the edge of the chip and the cells were lysed for 5 min.

If the prey protein was fluorescently tagged, the protein could be imaged immediately after cell lysis by using TIRF or confocal microscopy. Otherwise, 500 μL of a mixture containing 20 nM primary antibody against the prey protein and 30 nM fluorophore-labeled 2^nd^ antibody in T50-BSA buffer was added to the chip surface and incubated for 15 min, and after washing thrice with 1 mL of wash buffer to remove unbound antibodies, the prey protein on the chip could be examined under a confocal microscope. The entire procedure was performed at room temperature with the permanent magnet being used to hold the magnetic nanobeads at the bottom of the microwells.

### Single-molecule imaging and photobleaching under TIRF or confocal microscope

Microarray chips were examined using either an inverted TIRF microscope (IX73, Olympus) equipped with 488 and 561 nm lasers (OBIS, Coherent) or a confocal microscope (Leica TCS SP8). The procedures were the same as previously described ^6^, and all experiments were performed at room temperature.

### Data analysis and statistics

Signal intensity and signal-to-background (S/B) ratio were quantified as described ^6^. All data are expressed as means ± SD; n denotes the number of independent biological replicates. Unless indicated otherwise, Student’s two-tailed *t* test was used for statistical analysis, and *P* < 0.05 was considered statistically significant.

## Results

### Nanobead-based SiMPull strategy is simple and fast

The innovative SiMPull technique developed by Jain et al. enables detection of PPIs at the single-molecule level ^12^; however, wide application of this technique is hindered by its high technical barrier and time consumption. To overcome this hurdle, we previously used commercially available, pre-surface-functionalized agarose microbeads to develop a microbead-based SiMPull method ^5, 6^. The success of this work inspired us to further explore SiMPull based on magnetic nanobeads because magnetic nanobeads are adequately small to be drawn into microwells by applying magnetic force instead of using surface functionalization for single-cell analysis (see below). However, it was unclear how these tiny (∼100 nm) and opaque magnetic beads would work in SiMPull as compared to the larger (40–70 μm) and transparent agarose microbeads.

The nanobeads used in our method were also pre-blocked with a layer of BSA by the manufacturer to minimize nonspecific protein binding. Although the nanobeads are opaque, under a microscope, all fluorophores on a nanobead, whether facing the lens or not, can be excited and visualized because the nanobead diameter (∼100 nm) is smaller than visible-light wavelengths; therefore, these nanobeads are small “optically flat” surfaces. Similar to agarose-microbead-based SiMPull ^6^, our nanobead-based SiMPull (Fig. 1) saves time considerably by completely omitting the quartz-slide functionalization required in the method of Jain et al. ^12^, and our method is also highly reproducible because the surface-functionalized nanobeads used are subject to industry-level quality control.

We first characterized our nanobead-based SiMPull by using biotin-conjugated Alexa 488, which revealed that the sensitivity of the method was extremely high—the fluorophore could be detected at a concentration as low as 10 pM with a very high S/B ratio (>9) when IgG-conjugated Alexa 488 was used as a control (Supplementary Fig. 1). Moreover, we observed minimal nonspecific binding of proteins such as BSA and ovalbumin to the nanobeads (Supplementary Fig. 2) even after incubation for 4 h (Supplementary Fig. 3).

### Nanobead-based SiMPull for GFP pulldown from cell populations

Before using nanobead-based SiMPull for more demanding and challenging single-cell analyses, we validated the method for relatively simpler studies of cell populations, and we first assessed the performance of the method in the pulldown of GFP; this is because GFP can be directly visualized without immunostaining, which simplifies the validation, and, more importantly, because GFP can be used in photobleaching assays to determine the single-molecule state of a fluorescent spot captured on the nanobeads.

GFP ectopically expressed in HEK293T cells was efficiently pulled down by anti-GFP-coated magnetic nanobeads, but few GFP molecules were captured in 3 negative-control experiments (Fig. 2a); the calculated S/B ratio of the assay was ∼ 10–20 (Fig. 2b), which is comparable or superior to that of the original SiMPull method ^12^ or our microbead-based SiMPull ^6^. Furthermore, in photobleaching assays performed on the fluorescent spots by using our previously described procedures ^6^, we found that 58%, 18%, and 22% of the selected spots contained, respectively, single fluorophores (i.e., GFP monomers), 2 fluorophores, and >2 fluorophores (Fig. 2c-d). Conceivably, the proportion of single-fluorophore spots can be increased by diluting antibodies or GFP molecules ^6^. Nevertheless, our results clearly showed that the nanobead-based approach enables a protein of interest to be trapped at the single-molecule level at a very high S/B ratio, which represents one of the powerful features of the original ^12^ and microbead-based ^6^ SiMPull methods.

**Figure 2.**
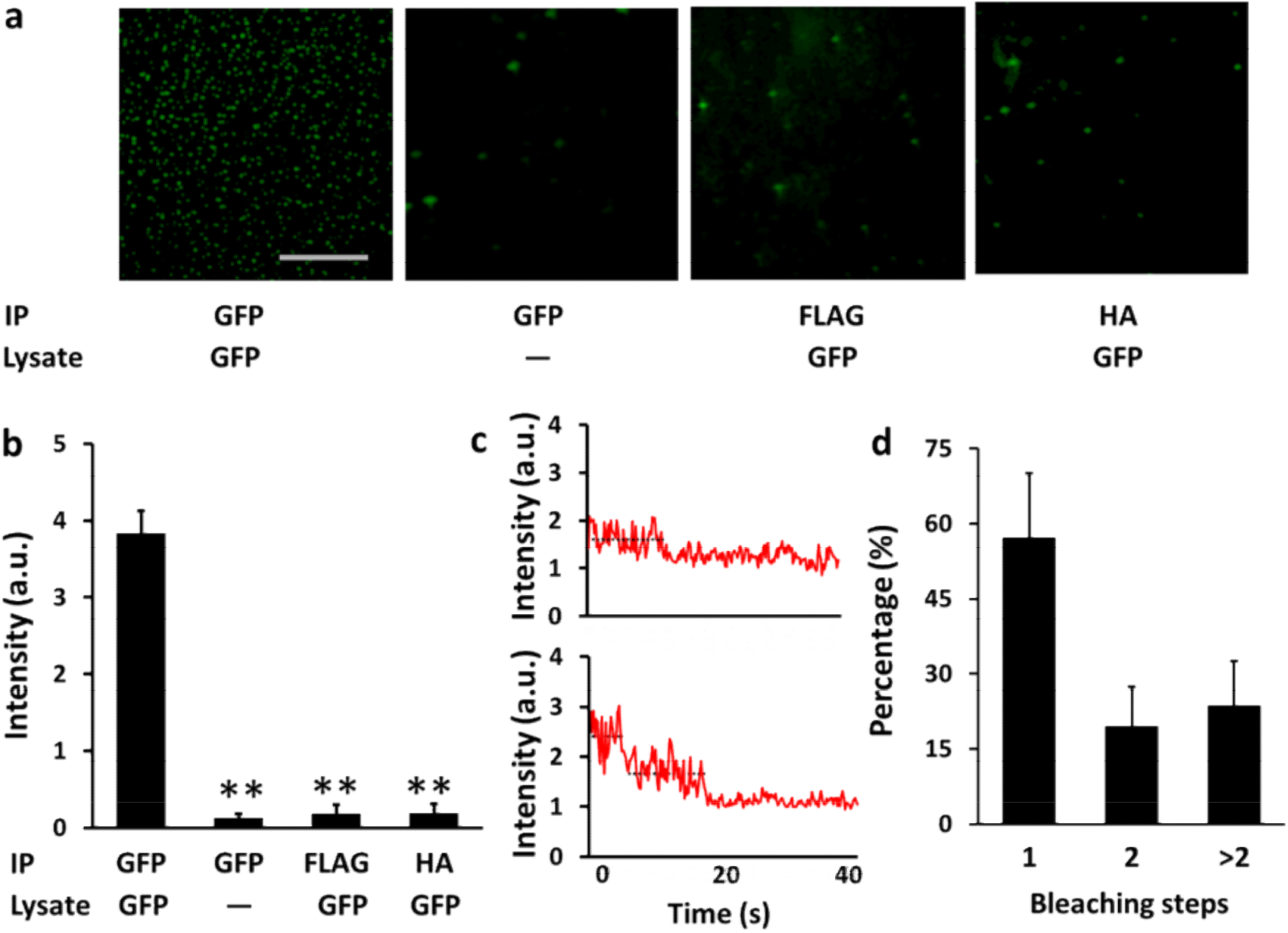
GFP pulldown from cell populations by using nanobead-based SiMPull. **a)** GFP was pulled down from pre-made lysates of GFP-expressing or non-transfected (control) HEK293T cells by using nanobeads sufaced-coated with anti-GFP, anti-FLAG, or anti-HA antibody and then examined under a TIRF microscope. The extremely high S/B ratio here suggests that GFP was spcifically pulled down by the anti-GFP antibody. The panel shows 30 × 30 μm imaging areas selected from a glass slide. Scale bar: 10 μm. **b)** Statistical results from experiment shown in **(a)** and 2 similar experiments. Different from 1^st^ group on the left: \*\**P* ≤ 0.0024; n = 3 independent biological replicates. **c-d)** Spots in the 1^st^ image on the left in **(a)** typically displayed one-step (upper) and two-step (lower) bleaching in photobleaching experiments **(c)**. Distribution of the photobleaching steps of 100 selected fluorescent spots in the image is shown in **(d)**. Most (∼60%) spots in an image typically displayed one-step bleaching in photobleaching experiments. n = 3 independent biological replicates; a.u., arbitrary unit.

### Pulldown of cAMP-dependent protein kinase A (PKA) complex from cell populations

The well-studied PKA holoenzyme or complex is a heterotetramer formed by a regulatory (R) subunit dimer and two catalytic (C) subunits; the binding of intracellular cAMP to the R subunits releases the C subunits from the PKA complex (Fig. 3a). We used this well-characterized interaction between the R and C subunits of PKA to evaluate the performance of the nanobead-based SiMPull assay in PPI analysis.

**Figure 3.**
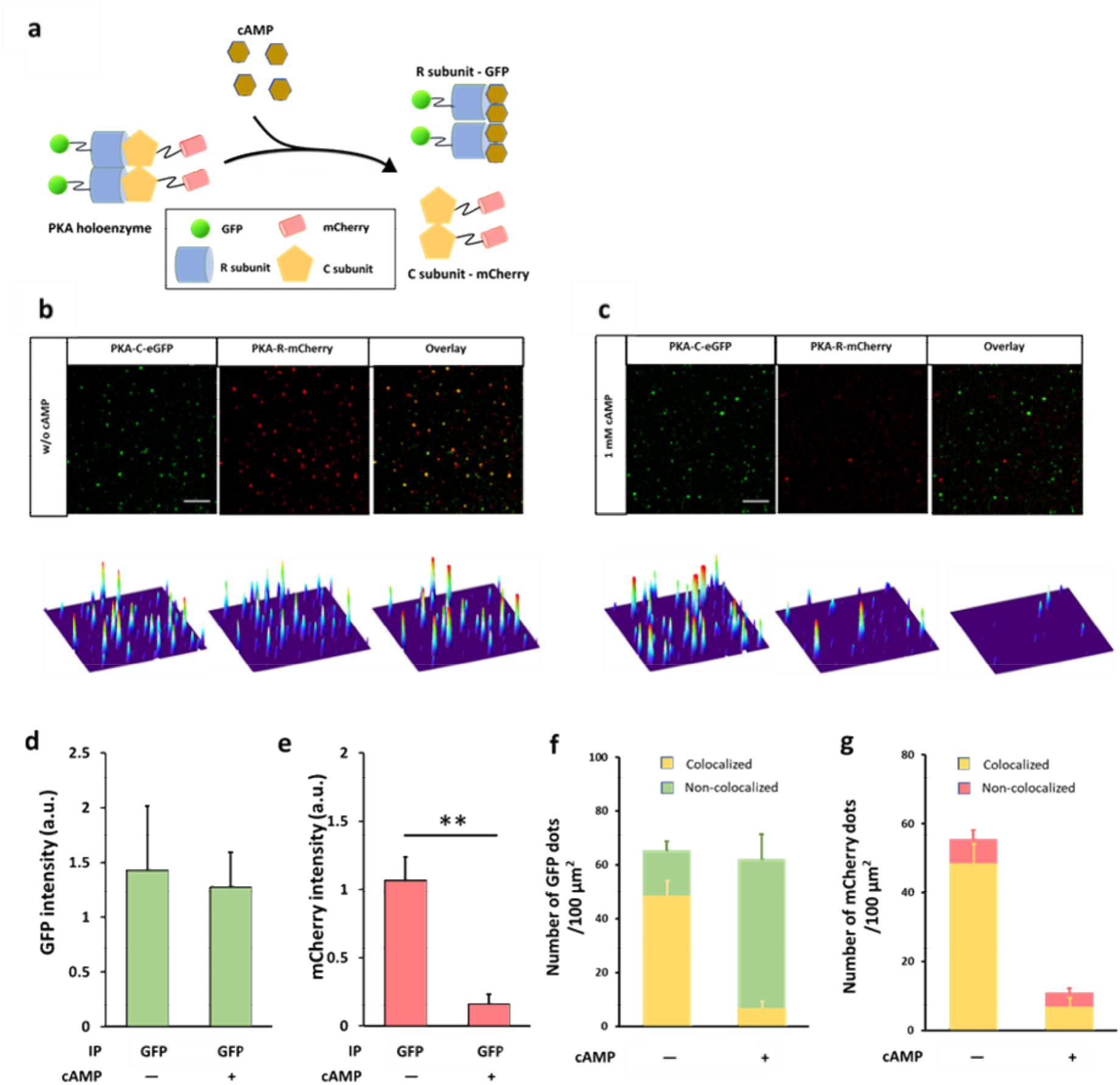
Pulldown of PKA complex from cell populations by using nanobead-based SiMPull. **a)** Schematic of PKA complex and its activation by cAMP. R, regulatory subunit; C, catalytic subunit. **b-c)** Upper images: Pulldown of PKA-C-eGFP (green) captured PKA-R-mCherry (red) in the absence of cpt-cAMP **(b)** but not in the presence of 1 mM cpt-cAMP **(c)**; lower images: corresponding surface plot maps of upper images. The panels show 18 × 18 μm imaging areas selected from a glass slide; scale bars: 3 μm. All analyses were performed using a confocal microscope. **d-e)** Statistical results of GFP **(d)** and mCherry **(e)** signal intensity in **(b-c)**. \*\**P* = 0.0096; n = 3 independent biological replicates, each representing the average of 3 imaging areas in the same experiment; a.u., arbitrary unit. **f-g)** Proportion of GFP **(f)** and mCherry **(g)** colocalized with each other in the absence and presence of 1 mM cpt-cAMP in **(b-c)**. The reduction in the amount of non-colocalized mCherry in the presence of cpt-cAMP suggests that some of the mCherry might be colocalized with quenched GFP in the absence of cpt-cAMP. n = 3 independent biological replicates used in **(d-e)**.

In the analysis, we examined the interaction between PKA-C-eGFP and PKA-R-mCherry in HEK293T cells because the fluorescent fusion proteins enable direct visualization of PKA-C and -R subunits without immunostaining and simplify the measurement of pulldown efficiency and specificity (Fig. 3b-e). When anti-GFP-coated nanobeads were used in the assay, both PKA-C-eGFP and PKA-R-mCherry were pulled down (Fig. 3b), which clearly indicated that the pulldown of PKA-C-eGFP co-immunoprecipitated its binding partner, PKA-R-mCherry. Furthermore, addition of the cAMP analog cpt-cAMP almost eliminated the pulldown of PKA-R-mCherry (Fig. 3c-e), which verified that the R subunit was captured because of its specific interaction with the C subunit rather than due to nonspecific binding to anti-GFP or the nanobeads. As expected, the majority of the GFP and mCherry signals clearly colocalized (Fig. 3f-g), but the colocalization was incomplete, which could be due to the several potential reasons discussed previously ^6^.

We also used GFP-fused PKA-C and -R subunits to perform photobleaching assays on the PKA complex captured on nanobeads (Supplementary Fig. 4): The selected fluorescent spots contained 4 fluorophores, which agrees with the stoichiometry of the heterotetrameric PKA complex, although in certain cases, the selected spots contained only 3 fluorophores, presumably because of the photobleaching of one of the subunits in the PKA tetramer or the release of one subunit from the complex (Supplementary Fig. 4). Nevertheless, these results clearly showed that our nanobead-based SiMPull enables a protein complex of interest to be trapped at the single-molecule level, which then allows analysis of the stoichiometry and binding kinetics of the protein complex ^12, 14^.

### Pulldown of endogenous protein complex from cell populations

Next, we validated nanobead-based SiMPull by examining the interaction between the endogenous soluble protein TRIP-Br1 and the transmembrane protein adenylyl cyclase 1 (AC1) in HeLa cells; the physical interaction between these two proteins has been well characterized previously ^13^. When TRIP-Br1 was immunoprecipitated from HeLa cell lysates, AC1 was also captured (Fig. 4a), but little AC1 signal was detected when either anti-AC1 was omitted in the immunostaining or anti-TRIP-Br1 was omitted in the immunoprecipitation (Fig. 4b-d). The calculated S/B ratio of the assay was ∼ 7.8–8.9. These results showed that our nanobead-based SiMPull technique can be successfully used to detect endogenous PPIs in cells.

**Figure 4.**
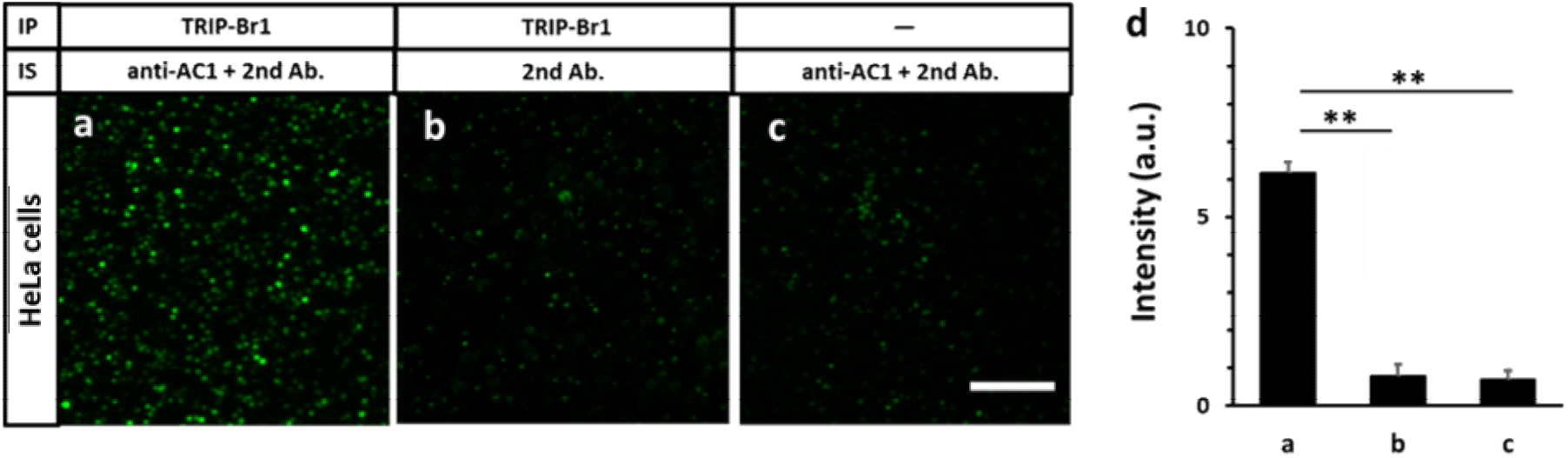
Pulldown of endogenous proteins from cell populations. **a-c)** Endogenous TRIP-Br1 was pulled down from 10^5^ HeLa cells by using magnetic nanobeads surface-coated with anti-TRIP-Br1 and then detected using anti-AC1 1^st^ antibody and fluorophore-labeled 2_nd_ antibody **(a)**. In negative-control experiments, AC1 was not detected in the absence of anti-AC1 **(b)** and was not pulled down by nanobeads that were not coated with anti-TRIP-Br1 **(c)**. The panels show 18 × 18 μm imaging areas selected from a glass slide; scale bar: 5 μm. **d)** Statistical results of experiments in **(a-c)**. *P* ≤ 0.0024; n = 3 independent biological replicates; S/B ratio: 7.8 (a/b) and 8.9 (a/c).

### Nanobead-based strategy for SiMPull from single cells

A key requirement for an optimal sc-SiMPull technique is a deep and narrow microchamber for trapping and lysing a single cell in situ; such a design can minimize cell-lysate diffusion and maximize interaction time between a target protein and its antibody for effective protein capture, and to achieve this, the surface of the microchamber bottom must be functionalized and coated with antibodies. However, functionalizing a large surface is cumbersome, as discussed previously ^6^, and functionalizing a microscale surface is expected to be considerably more challenging because the deep and narrow microchamber makes the multiple solution exchanges required in the original procedure ^12^ extremely slow, time-consuming, and irreproducible. This technical challenge might explain why a cell-trapping microchamber was not used in two previously reported sc-SiMPull methods ^9, 10^, an omission that substantially compromises the applicability of the methods (see “Discussion”). Our nanobead-based SiMPull method (Figs. 1–4) provides an innovative and highly effective solution to this technical bottleneck: placing pre-surface-functionalized magnetic nanobeads on the glass bottom of microwells (Fig. 5) instead of directly functionalizing the glass bottom.

**Figure 5.**
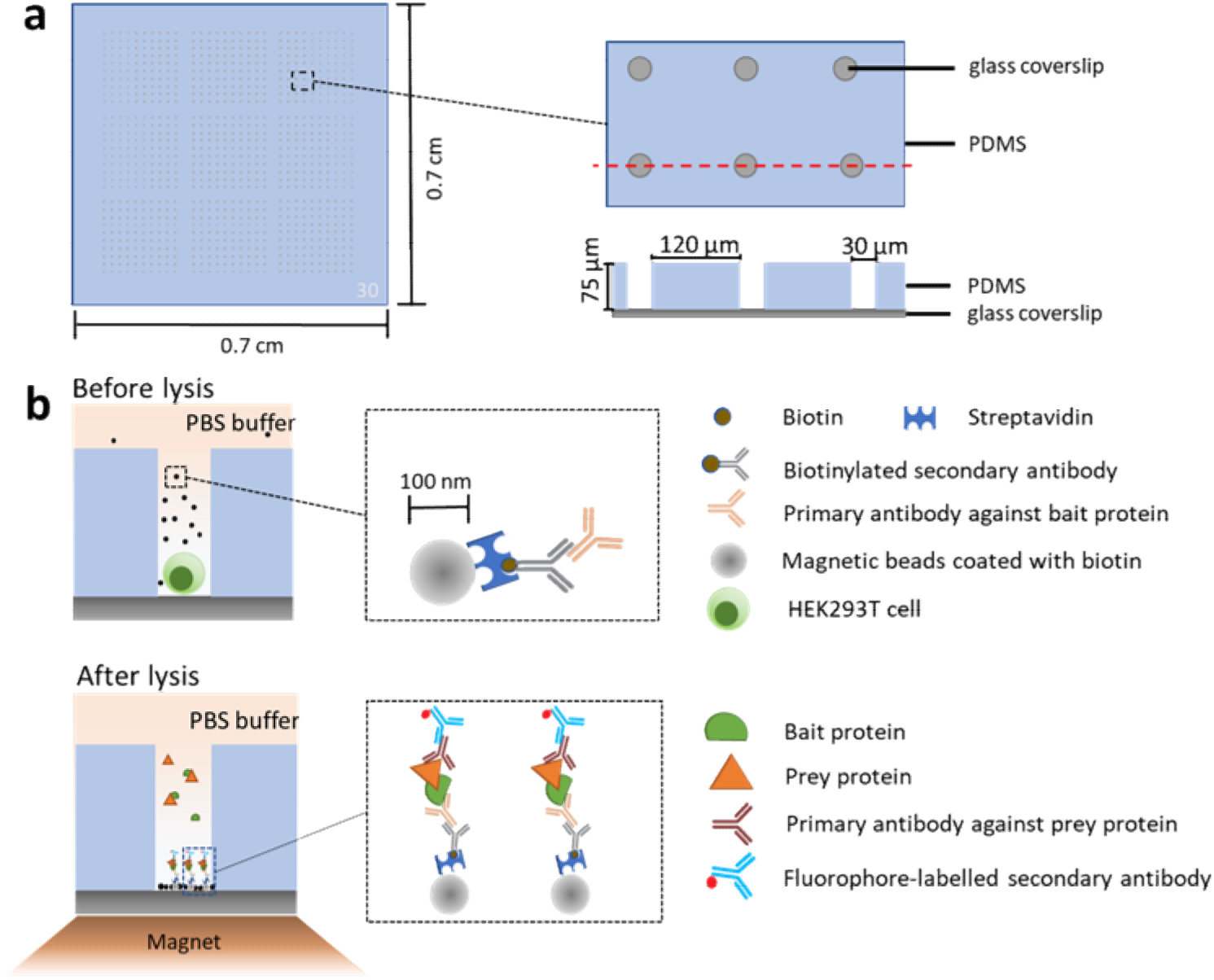
Schematic of nanobead-based SiMPull for single cells. **a)** Microwell array chip: schematic top view (whole, left; partial, upper right) and cross-section through dotted red line (lower right). Each microwell is 30 μm wide and 70 μm deep. **b)** Upper (before lysis): After a single cell is trapped in a microwell, surface-modified magnetic nanobeads (70–130 nm in diameter) are applied to the microwell chip. Streptavidin-coated magnetic nanobeads are modified by immobilizing on their surface a biotinylated 2^nd^ antibody and a specific 1^st^ antibody for capturing the target (bait) protein. Lower (after lysis): To lyse captured cells, a lysis buffer is added to the top of the chip. After cell lysis and protein capture, magnetic nanobeads are dragged to the microwell bottom by a magnet and the prey protein is detected using a specific 1^st^ antibody and a fluorescently labeled 2^nd^ antibody. The nanobeads are washed thrice and imaged using TIRF or confocal microscopy. If the bait protein or prey protein is fluorescently labeled, the protein can be visualized directly without immunostaining.

The microwell array chip here was prepared following a previously published procedure ^15^ with certain modifications (Fig. 5a and Supplementary Fig. 5). The microwell diameter was 30 μm but adjustable to fit the target-cell size (range 15–50 μm), and the microwell was designed to be deep (70 μm) to effectively trap single cells and minimize cell-lysate diffusion (Fig. 5a). In our experiments, ∼40% of the total microwells in a chip were occupied by cells and ∼85% of cell-occupied microwells contained single cells (Supplementary Fig. 6).

After single cells were trapped in microwells, magnetic nanobeads pre-coated with bait antibodies were applied. In the absence of a magnet, the nanobeads sedimented extremely slowly, but when a magnetic force was applied, the nanobeads were pulled down to the microwell bottom in 3 min (Fig. 5b and Supplementary Fig. 7). Subsequently, the lysis buffer was added to the chip to lyse the cells in situ; the bait antibody immobilized on the nanobeads captured the bait and prey proteins present in the lysate within the microwell; and the magnetic nanobeads were immobilized at the glass bottom of the microwells by the magnet during the entire experiment. Lastly, the prey proteins, either fluorescently tagged or labeled through immunostaining, were visualized and identified using TIRF or confocal microscopy (Fig. 5b).

### GFP and ANO1 pulldown from single cells

As with the experiments shown in Fig. 2, we first validated the nanobead-based sc-SiMPull assay by pulling down GFP from single cells. After suspension through trypsinization, GFP-expressing HEK293T single cells were added to a blank microwell chip (Fig. 6a-b) and anti-GFP-coated magnetic nanobeads were added (Fig. 6c and Supplementary Fig. 7), and the single cells were then lysed and the magnetic nanobeads were used for the pulldown. Whereas GFP was pulled down by nanobeads coated with anti-GFP through a biotinylated 2^nd^ antibody (Fig. 6d-e and Supplementary Video 1), GFP was not captured by nanobeads coated with the 2^nd^ antibody alone (Fig. 6f-g). Thus, GFP was captured because of its specific interaction with anti-GFP rather than due to nonspecific binding to the 2^nd^ antibody or the nanobeads (Fig. 6h); the S/B ratio in these experiments was ∼12 (Fig. 6h). Moreover, photobleaching assays revealed that ∼40% of the GFP fluorescent spots displayed one-step bleaching (Fig. 6i-j), which indicated the presence of GFP monomers.

**Figure 6.**
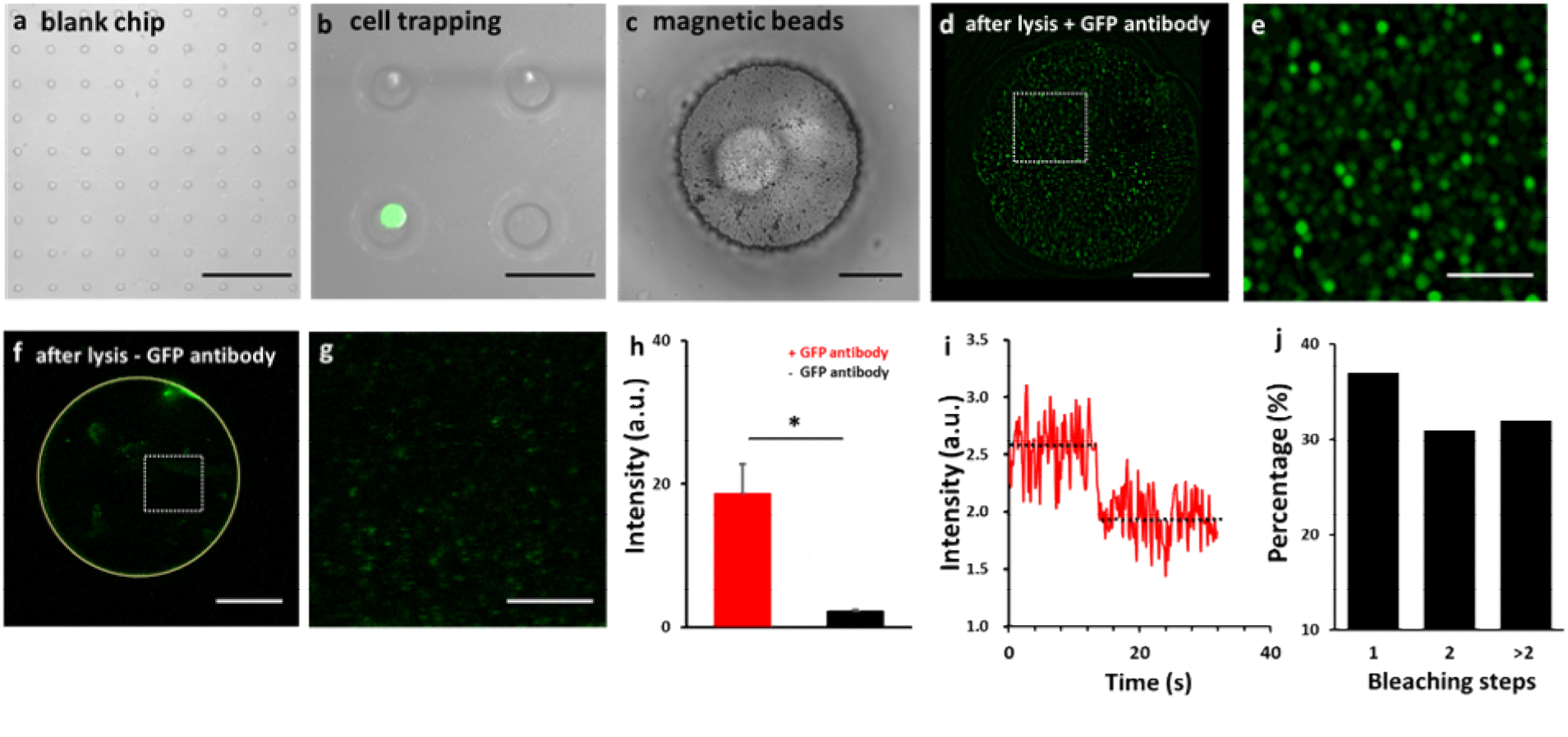
GFP pulldown from single cells. **a)** Top view of part of a blank microwell chip (each microwell is 30 μm in diameter). **b)** Top view of 4 microwells, one of which (lower left) has trapped a single GFP-expressing HEK293T cell (green). **c)** Top view of a microwell containing a GFP-expressing HEK293T cell after application of magnetic nanobeads (small black particles) to the chip. **d-e)** Single microwell showing fluorescence of GFP pulled down by anti-GFP-coated magnetic nanobeads after cell lysis **(d)**, and magnification of boxed area **(e)**. **f-g)** Single microwell (outlined by yellow line) in a negative-control experiment **(f)**, and magnification of boxed area **(g)**. GFP was not pulled down by nanobeads coated with 2^nd^ antibody but not anti-GFP. A certain amount of nonspecific signal was observed at the microwell edge. **h)** Total intensity of fluorescent dots in **(d)** and **(f)** and two independent biological replicates. \**P* = 0.033. **i-j)** A GFP spot in **(e)** displayed one-step photobleaching in a photobleaching assay. Distribution of the photobleaching steps of GFP molecules in **(d)** is shown in **(j)**. All experiments shown in this figure were performed using a TIRF microscope. a.u., arbitrary unit; scale bars (in μm): 300 in **(a)**; 50 in **(b)**; 10 in **(c)**, **(d)**, and **(f)**; 3 in **(e)** and **(g)**.

As compared to analyzing soluble proteins such as GFP, studying transmembrane proteins in pulldown assays poses an additional challenge: the transmembrane proteins must first be well solubilized, and unsolubilized membrane debris can potentially interfere with the pulldown assays, particularly SiMPull assays employing microscopy for detection. In cell-population analyses, this technical hurdle can be readily overcome by removing the unsolubilized membrane debris through centrifugation; however, it was unclear how this problem would affect the performance of nanobead-based sc-SiMPull, where single cells are lysed in a microwell in situ and any unsolubilized membrane debris would remain in the microwell. To assess the performance of nanobead-based sc-SiMPull in the study of transmembrane proteins, we tested the pulldown of ANO1(TMEM16A)-GFP from HEK293T single cells. ANO1-GFP was pulled down by anti-GFP-coated magnetic nanobeads but not by nanobeads without the anti-GFP coating (Supplementary Fig. 8); the calculated S/B ratio here was 6.4 (Supplementary Fig. 8b). However, the homogeneity of the fluorescent spots was not as high as that in the experiments involving soluble proteins (Figs. 6–7), and this could result from incomplete solubilization of ANO1-GFP. Nevertheless, our results showed for the first time that sc-SiMPull can be used for pulling down transmembrane proteins with an acceptably high S/B ratio.

**Figure 7.**
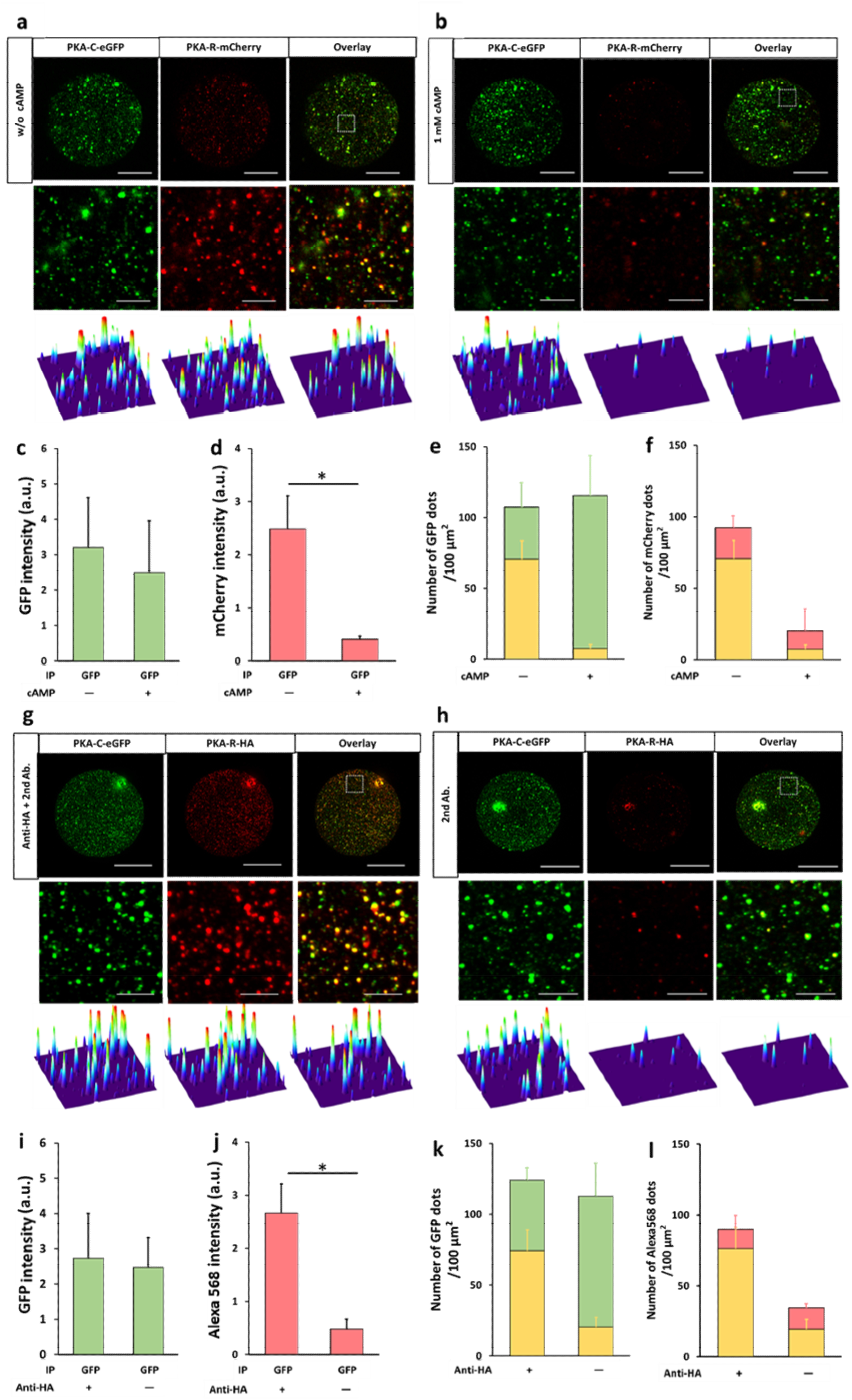
Pulldown of PKA complex from single cells. **a-b)** Anti-GFP-coated magnetic nanobeads pulled down both PKA-C-eGFP (green) and PKA-R-mCherry (red) from single cells in the absence of cAMP **(a)** but pulled down only PKA-C-eGFP (green) in the presence of cAMP **(b)**. In **(a-b)** (and also in **(g-f)**), middle and bottom images are magnified views of boxed areas in upper images and their corresponding surface plot maps, respectively; scale bars: upper, 20 μm; lower, 3 μm. **c-f)** Statistical results of signal intensity in **(a-b)** are shown in **(c-d)**, respectively, and **(e-f)** show proportions of GFP **(e)** and mCherry **(f)** colocalized with each other in **(a-b)**; the reduction in the amount of non-colocalized mCherry in the presence of cpt-cAMP suggests that some of the mCherry might be colocalized with quenched GFP in the absence of cpt-cAMP **(b)**. n = 3 independent biological replicates, each representing the average of 2 images of an entire well in the same experiment; \**P* = 0.0311. **g-h)** Anti-GFP-coated magnetic nanobeads pulled down PKA-C-eGFP (green) and PKA-R-HA (red) from single cells. PKA-R-HA was visualized by immunostaining with anti-HA and Alexa 568-conjugated 2^nd^ antibody **(g)**. Few HA-positive spots were detected in immunostaining in the absence of anti-HA **(h)**, suggesting that the HA staining observed in **(c)** was HA-specific and not nonspecific binding of the 2^nd^ antibody. **i-l)** Statistical results of signal intensity in **(g-h)** are shown in **(i-j)**, respectively, and **(k-l)** show the proportions of GFP **(k)** and anti-HA (Alexa 568) **(l)** colocalized with each other in **(g-h)**. n = 3 independent biological replicates, each representing the average of 2 images of an entire well in the same experiment; \**P* = 0.0304.

### Pulldown of PKA complex from single cells

Lastly, we again used the well-characterized interaction between PKA-R and -C subunits to validate nanobead-based sc-SiMPull for studying PPIs. We examined the interaction between PKA-C-eGFP and PKA-R-mCherry in single HEK293T cells (Fig. 7a-f). Similar to the results shown in Fig. 3, anti-GFP-coated nanobeads pulled down both PKA-C-eGFP and PKA-R-mCherry under control conditions (Fig. 7a), but few PKA-R-mCherry molecules were captured in the presence of cpt-cAMP (Fig. 7b). Moreover, the majority (∼70%) of the GFP and mCherry signals clearly colocalized (Fig. 7g-h).

We also evaluated the nanobead-based sc-SiMPull assay for more general application by using a prey protein, PKA-R, that was not fluorescently tagged and thus required immunostaining for visualization. Here, single HEK293T cells coexpressing PKA-C-eGFP and PKA-R-HA were subject to sc-SiMPull by using anti-GFP-coated nanobeads, and after cell lysis and protein capture, a mixture of anti-HA and Alexa 561-conjugated 2^nd^ antibody was added to the chip surface to visualize PKA-R-HA. PKA-R-HA was clearly visible after staining with anti-HA but was not detected if anti-HA was omitted (control) (Fig. 7g-h), and ∼ 60% of the GFP spots colocalized with the Alexa 561 spots (Fig. 7k). The colocalization ratio here was lower than that in the experiments shown in Fig. 3f or Fig. 7e probably because the immunostaining shown in Fig. 7g required the binding of 1^st^ and 2^nd^ antibodies and therefore likely did not detect all PKA-R molecules.

## Discussion

The original SiMPull assay for cell populations developed by Jain and colleagues is ultrasensitive and highly versatile and has substantially increased our capacity to analyze PPIs ^12^. However, wide application of this method ^12^ and its high-throughput variation ^16^ is considerably impeded by their high technical barrier and time consumption (Zhao et al. 2021). To overcome these challenges, we previously developed an agarose-microbead-based SiMPull assay for cell populations ^6^. Relative to the original SiMPull method, our microbead-based SiMPull is considerably faster, easier to use, and more reproducible, and yet provides similar sensitivity and S/B ratio ^5, 6^.

In this study, we expanded the bead-based approach by examining the performance of a SiMPull assay based on magnetic nanobeads for cell populations; our ultimate aim here was to use nanobead-based SiMPull for single-cell analysis if the method performed well in the cell-population analyses. Notably, in cell-population SiMPull, the performance of magnetic nanobeads was similar to that of agarose microbeads in terms of assay simplicity, time consumption, and S/B ratio (Figs. 1–4). Thus, we next evaluated nanobead-based SiMPull for single-cell analysis (Fig. 5) and found that the method enabled the pulldown of both a soluble protein (GFP) and a transmembrane protein (ANO1) with a very high S/B ratio (Fig. 6 and Supplementary Fig. 8); moreover, nanobead-based sc-SiMPull captured protein complexes such as the PKA holoenzyme and allowed determination of the stoichiometry of the complex in single cells (Fig. 7 and Supplementary Fig. 4).

As compared to the previously published sc-SiMPull methods ^9–11^, which are mostly variations of the original SiMPull approach ^12^, our nanobead-based sc-SiMPull method presents a considerably lower technical barrier. Furthermore, our method offers several other advantages. Because of the lack of a cell-trapping microchamber to minimize cell-lysate diffusion, two of the previously reported sc-SiMPull techniques can be used for analyzing slowly diffusing and soluble molecules only in bacteria ^10^ or adherent-cell cultures ^9^ and not in primary- or suspension-culture cells (blood cells, circulating tumor cells, etc.). Moreover, in the Wedeking et al. method ^9^, cells are cultured on strips of cell-adhesion molecules; this artificial culture condition involves the use of complex fabrication procedures and can generate artifacts in terms of the mechanical and chemical properties of cells.

The lack of a cell-trapping microchamber in the aforementioned two methods also leads to the requirement of slow release of cytosolic proteins to allow sufficient time for proteins to be captured ^9, 10^. To ensure this slow protein release, the extraction step must avoid complete cell lysis by using strong detergents, which are however essential for adequately solubilizing transmembrane proteins. By contrast, our sc-SiMPull can be used for pulling down transmembrane proteins (Supplementary Fig. 8) because a physical diffusion barrier is formed by the microwell in our method and therefore the stronger solubilization conditions necessary for capturing transmembrane proteins can be accommodated. Relative to soluble proteins, transmembrane proteins are typically present in substantially lower amounts in the cell and more frequently require ultrasensitive techniques such as SiMPull for analysis.

Cell-trapping is possible in one previously reported sc-SiMPull assay ^11^, but the design of the cell-trapping chamber here is complex and can accommodate only one single cell (zygote of *Caenorhabditis elegans*). The high technical barrier and extremely low throughput in this case clearly limit the application of this method, which has also not been tested thus far for the pulldown of any transmembrane protein.

In summary, we have developed a new sc-SiMPull strategy that can, in principle, be used for studying PPIs or protein-DNA/RNA interactions. Relative to previously reported sc-SiMPull methods, our nanobead-based approach is considerably faster, easier to use, more reproducible, more versatile for distinct cell types and protein molecules, and yet provides similar sensitivity and S/B ratio.

## ACKNOWLEDGMENTS

The work was supported by Hong Kong RGC GRF16102417, GRF16100218, GRF16102720, and GRF16101321, NSFC-RGC Joint Research Scheme N_HKUST614/18, SMSEGL20SC01-K (all to P.H.), and in part by the Innovation and Technology Commission (ITCPD/17-9).

## AUTHOR CONTRIBUTIONS

Conceptualization: Q.Z., Y.S., and P.H.; methodology: Q.Z., Y.S., F.T., L.Y., and H.P.; formal analysis: Q.Z. and P.H.; investigation: Q.Z., X.L., X.Y., Y.L., and Z.L.; resources: R.T. and H.P.; writing-original and draft: Q.Z. and P.H.; writing-review and editing: P.H. with contributions from all the authors; visualization: Q.Z.; supervision: P.H.; funding acquisition: P.H.

## DATA AVAILABILITY STATEMENT

The data that support the findings of this study are available on request from the corresponding author, [Pingbo HUANG], upon reasonable request.

